# The role of sodium channels in sudden unexpected death in pediatrics

**DOI:** 10.1101/383562

**Authors:** Anne M. Rochtus, Richard D. Goldstein, Ingrid A. Holm, Catherine A. Brownstein, Eduardo Pérez-Palma, Dennis Lal, Annapurna H. Poduri

## Abstract

Sudden Unexpected Death in Pediatrics (SUDP) is a tragic condition with hypothesized multifactorial etiology. While there is recent evidence implicating genes related to cardiac arrhythmia and epilepsy as genetic risk factors contributing to some cases of SUDP, the underlying mechanisms of SUDP remain under active investigation. SUDP encompasses Sudden Infant Death Syndrome (SIDS) and Sudden Unexplained Death in Childhood (SUDC), affecting children under and over 1 year of age, respectively. The presence of developmental hippocampal malformations in many children with SIDS and SUDC suggests that a subset of patients may share epilepsy-related mechanisms with Sudden Unexplained Death in Epilepsy Patients (SUDEP). Pathogenic variants in both epilepsy- and arrhythmia-related sodium channel genes have recently been identified in patients with SIDS, SUDC, and SUDEP.

We performed a candidate gene analysis for genes encoding sodium channel subunits in whole exome sequencing (WES) data from 73 SUDP patients. After a thorough literature review, we mapped all reported SUDP-associated sodium channel variants alongside variants from the population on a structural protein model to evaluate whether patient variants clustered in important protein domains compared to controls.

In our cohort, 13 variants met criteria for pathogenicity or potential pathogenicity. While *SCN1A, SCN1B*, and *SCN5A* have established disease associations, we also considered variants in the paralogs *SCN3A, SCN4A* and *SCN9A*. Overall, the patient-associated variants clustered at conserved amino acid sites across the sodium channel gene family that do not tolerate variation in these genes.

This study provides a molecular overview of sodium channel variants present in cases with SUDP and reveals key amino acid sites that do not tolerate variation across the SCN paralog family. Further research will lead to an improved understanding of the contribution of sodium channels to SUDP, with a goal of one day implementing prevention strategies to avoid untimely deaths in at-risk children.

**Author Summary:** The sudden unexplained death of an infant or a child is a tragic event, which is likely caused by the complex interaction of multiple factors. Besides environmental factors, genes related to epilepsy and cardiac arrhythmia have been identified as risk factors. The sodium channel family encompasses genes, related to both cardiac arrhythmia as well as epilepsy, whose proteins share structural homology. We evaluated sodium channel gene variants in our cohort, examined all known variants in sodium genes in SUDP patients from the literature, and mapped patient variants alongside variants from the population on a 3D protein model. The patient variants clustered at conserved amino acid sites with low rates of variation in the general population, not only in the particular gene involved but also in the gene family. This study illustrates that sodium channel variants contribute to the complex phenotype of sudden death in pediatrics, suggesting complex mechanisms of neurologic and/or cardiac dysfunction contributing to death.

## Introduction

Sudden Unexpected Death in Pediatrics (SUDP) encompasses a tragic set of conditions, including Sudden Infant Death Syndrome (SIDS) and Sudden Unexplained Death in Childhood (SUDC), affecting children under and over 1 year of age, respectively. These conditions are hypothesized to involve heterogeneous and multifactorial etiologies, conceptualized as a ‘triple-risk’ model with a convergence of intrinsic, developmental, and environmental vulnerabilities contributing to death [1,2]. We have reported developmental hippocampal malformations in greater than 40% of children with SIDS and SUDC [3,4], suggesting that a subset of SIDS and SUDC is linked to epilepsy-related mechanisms since such hippocampal lesions have been classically associated with temporal lobe epilepsy [5]. The association between epilepsy and sudden death, demonstrated most clearly in sudden unexpected death in epilepsy (SUDEP), may well extend to SIDS and SUDC in patients with these lesions, which have been called ‘epilepsy *in situ*’ [6]. While the terminal mechanisms of SUDP and SUDEP remain speculative [7,8], there is active investigation into the role of genetic factors involving genes related to epilepsy [9,10] as well as cardiac arrhythmias [11,12].

Voltage-gated sodium channels (VGSCs) are a highly conserved family of proteins—expressed in excitable tissue in heart, central nervous system, peripheral nervous system, and muscle—that are essential for the generation and propagation of action potentials. Interestingly, pathogenic variants in both arrhythmia- and epilepsy-related VGSCs have been identified in patients with SIDS, SUDC, and SUDEP [9,12-30]. In humans, nine different pore-forming α-subunits have been identified (Na_v_ 1.1–1.9 encoding for *SCN1A-SCN5A* and *SCN8A-SCN11A*) [31,32]. Na_v_ 1.1, 1.2, 1.3 and 1.6 are the primary sodium channel subunits expressed in the central nervous system, Na_v_ 1.7, 1.8 and 1.9 in the peripheral nervous system, Na_v_ 1.4 in skeletal muscle, and Na_v_ 1.5 in the heart. The pore-forming α-subunit is composed of four homologous domains, each containing six transmembrane α-helical segments (S1-S6). In addition, there are five different β-subunits (β1, β1B, β2, β3, β4) encoded by *SCN1B-SCN4B* [33], The tissue-specific expression profiles of α-subunits and β-subunits are shown in Table 1. Variants in the cardiac-expressed *SCN5A* [12-26] gene are reported most frequently in association with SIDS and SUDC, but variants in *SCN1A* [9,27], *SCN4A* [34], *SCN10A* [35], *SCN1B* [12,35-38], *SCN3B* [15,39] and *SCN4B* [39] have also been reported. In addition, variants in *SCN1A* [40,41], *SCN2A* [28,42] and *SCN8A* [29,30,42] have been associated with SUDEP.

**Table 1.** Voltage-gated sodium channels expression and disease associations.

| Gene | Protein | Distribution | Associated human disease |
| --- | --- | --- | --- |
| ***SCN1A*** | Nav1.1 | CNS, heart | DS, GEFS+, FS+, familial autism, FHM3, SUDEP |
| ***SCN2A*** | Nav1.2 | CNS | DS, GEFS+, OS, EOEE, BFNIS |
| ***SCN3A*** | Nav1.3 | CNS, heart | Unclear, contributing to neuronal hyperexcitability/ epilepsy? |
| ***SCN4A*** | Nav1.4 | Skeletal muscle | PAM, PMC, HyperPP, HypoPP, SNEL |
| ***SCN5A*** | Nav1.5 | Skeletal muscle, heart, CNS | AF, AS, BS, DCM, LQTS, PCCD, SIDS, SSS, SUDEP |
| ***SCN8A*** | Nav1.6 | CNS, PNS | EOEE, cognitive impairment, paralysis, ataxia, dystonia |
| ***SCN9A*** | Nav1.7 | PNS | CIP, IEM, PEPD, PPN |
| ***SCN10A*** | Nav1.8 | PNS | PPN |
| ***SCN11A*** | Nav1.9 | PNS | PPN |
| ***SCN1B*** | $\beta 1$ | CNS, PNS, heart | AF, BS, DS, GEFS+, LQTS, PCCD, TLE |
| ***SCN2B*** | $\beta 2$ | CNS, PNS, heart | AF, BS |
| ***SCN3B*** | $\beta 3$ | CNS, PNS, heart | AF, BS, PCCD, SIDS, ventricular fibrillation |
| ***SCN4B*** | $\beta 4$ | CNS, PNS, heart | LQTS, SIDS |
| ***SCN1B*** | $\beta 1B$ | Fetal CNS, PNS, heart | BS, PCCD, epilepsy |

Legend: Modified with permission from Brunklaus et al [43].
AF, Atrial fibrillation; AS, atrial standstill; BFNIS, benign familial neonatal-infantile seizures; BS, Brugada syndrome; CIP, channelopathy-associated insensitivity to pain; CNS, central nervous system; DCM, dilated cardiomyopathy; DS, Dravet syndrome; EOEE, early-onset epileptic encephalopathy; FHM3, familial hemiplegic migraine type 3; FS+, febrile seizures plus; GEFS+, genetic epilepsy with febrile seizures plus; HyperPP, hyperkalemic periodic paralysis, HypoPP, hypokalemic periodic paralysis; IEM, inherited erythromelalgia; LQTS, long QT syndrome; OS, Ohtahara syndrome; PAM, potassium-aggravated myotonia; PCCD, progressive cardiac conduction disease; PEPD, paroxysmal extreme pain disorder formally known as familial rectal pain syndrome; PMC, paramyotonia congenital; PNS, peripheral nervous system; PPN, painful peripheral neuropathies; SIDS, sudden infant death syndrome; SNEL, severe neonatal episodic laryngospasm; SSS, sick sinus syndrome; SUDEP, sudden unexplained death in epilepsy; TLE, temporal lobe epilepsy.

Given the common evolutionary origin and expression pattern of sodium channel subunits expressed across cardiac and neurologic tissues, we performed a candidate gene analysis for variants in genes encoding these proteins and their paralogs using whole exome sequencing (WES) data from 73 patients with SUDP. In addition, we performed a structure-based assessment of all novel and reported variants in human sodium channels in patients with SUDP vs. controls.

## Results

### Genetics and clinical characteristics of our cases

We identified 13 variants that we determined to be pathogenic or likely pathogenic in genes encoding for VGSCs in 11 patients, using ACMG criteria [44] (Table 2). The age of death across the 11 patients with variants in VGSC-encoding genes ranged from 7 weeks to 8 years, with 9/11 (81%) patients younger than 6 months at the time of death. Eight patients had hippocampal malformations as assessed by detailed neuropathological examination, and 1 patient had a normal hippocampus. For 2 other patients, detailed neuropathological analysis was not possible. Full clinical and molecular data for all 11 patients are listed in Table 2. Notably, one patient had two variants in *SCN1A* (p.Leu1296Met, p.Glu1308Asp) (reported previously)[9], one patient had a variant in *SCN3A* (p.Ala1804Val) and in *SCN10A* (c.4386+1G>C), and two siblings carried the same variant in *SCN1B* (p.Trp179Ter).

**Table 2.**
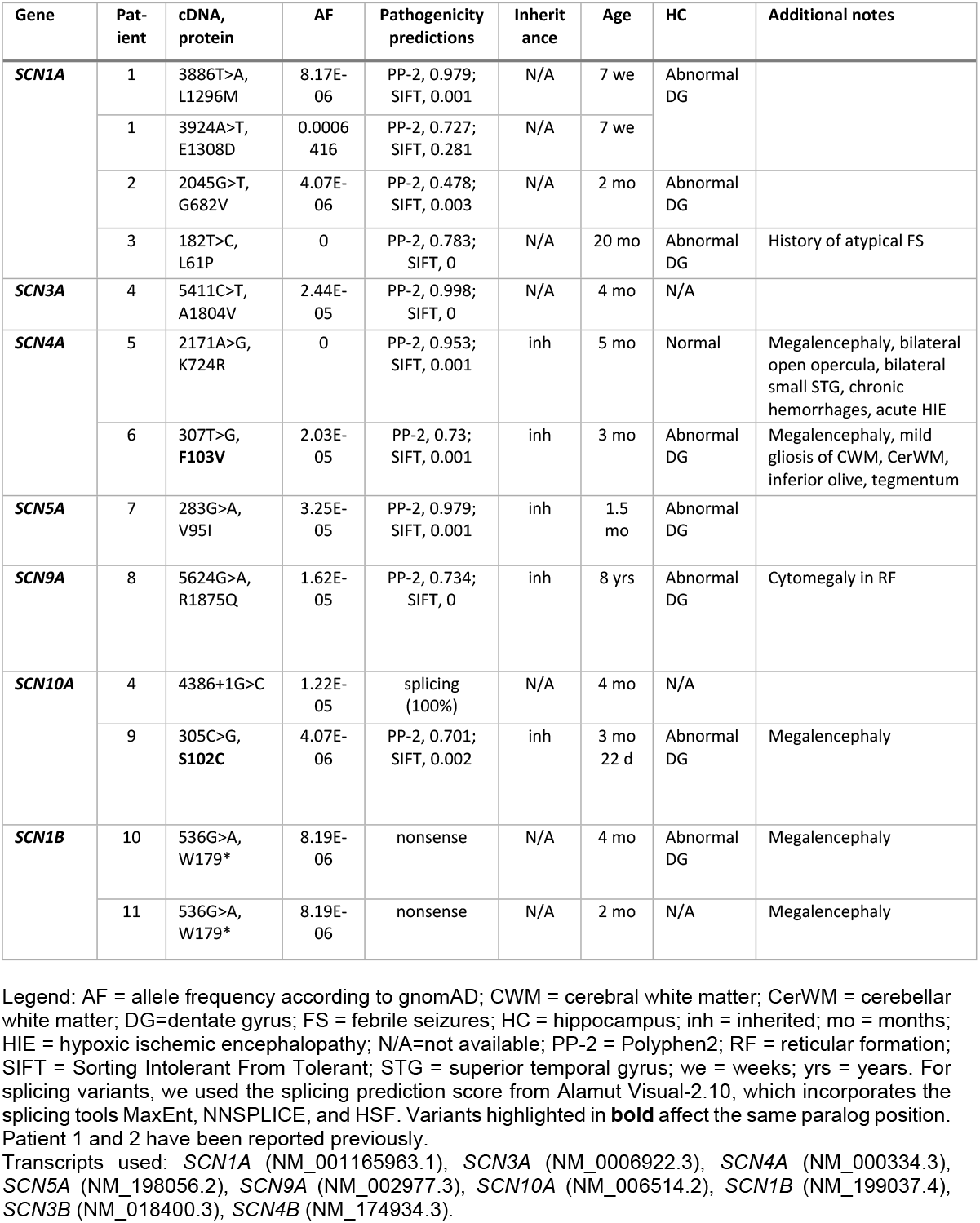
Variants in voltage-gated sodium channel genes in our cohort.

### Cases from the literature

We reviewed all variants in VGSC genes that have been reported in the literature in patients with SIDS and SUDC. We identified 74 variants in 99 patients affecting 72 different amino acid positions in the following genes: *SCN1A* (n=3), *SCN4A* (n=5), *SCN5A* (n=55), *SCN10A* (n=4), *SCN1B* (n=3), *SCN3B* (n=3), *SCN4B* (n=1).

### Re-evaluation of all variants

Collectively, 13 patients’ variants from our own cohort plus 99 patients’ variants retrieved from literature comprise 84 variants affecting 82 different amino acid positions in 9 genes: *SCN1A* (n=4), *SCN3A* (n=1), *SCN4A* (n=8), *SCN5A* (n=56), *SCN9A* (n=1), *SCN10A* (n=6), *SCN1B* (n=4), *SCN3B* (n=3), *SCN4B* (n=1). Three of the variants included here were previously reported [9,27].

Evaluating variants in the literature with respect to allele frequency reported in the general population, 11/74 variants (15%) of the variants reported in the literature had an allele frequency higher than 0.001, arguing against their pathogenicity. In addition, using pathogenicity prediction scores, we identified 23 additional variants that were predicted to have a benign or neutral functional effect. In total, we considered that there was conflicting evidence of pathogenicity for 33 out of the 74 (45%) variants reported in literature, and 1 out of 74 (1%) was determined to be benign based on the high frequency in controls, lack of predicted functional effect *in silico*, and/or *in vivo* absence of functional effects resulting from the variants (S1 Table).

To further assess for evidence of variant pathogenicity, we determined the Parazscore for all missense variants. As expected, Parazscores from patient variants were significantly higher than those observed in gnomAD controls (p-value < 0.0001). The patients’ variants that we determined to be pathogenic with the prediction tools were more likely to be located at strongly conserved family members (p=0.03) (i.e., greater Parazscores) (Fig 1A). On the other hand, patients’ variants that we determined to be conflicting or benign with the prediction scores were more likely present in less conserved regions (p=0.0001) (Fig 1B). Interestingly, 25 variants involved the ‘alignment index position’ of a VGSC variant reported in disease (S2 Table), which is unlikely to occur in the general population (p-value < 0.0001). Overall, we observed significant clustering of variants at conserved amino acid sites, notably with the same amino acid affected between *SCN1A/SCN5A* and *SCN5A/SCN9A* (S2 Table). The variants of two patients (Patients 6 and 9) in our cohort affected the same paralog position: *SCN4A* (p.Phe103Val) and *SCN10A* (p.Ser102Cys) (p-value < 0.0001). Both patients died at 3 months of age and had pathology notable for hippocampal granule cell dispersion with dentate gyrus bilamination (Table 2); both variants were inherited from a parent who had no history of epilepsy, febrile seizures, or other major illness. Three other patients of our cohort (Patients 1, 7 and 8) affected the same paralog position at respectively *SCN1A/SCN5A, SCN5A/SCN9A*, and *SCN5A/SCN10A*. Interestingly, all three patients had dentate gyrus bilamination (Table 2 and Supplementary Table 2).

**Fig 1:**
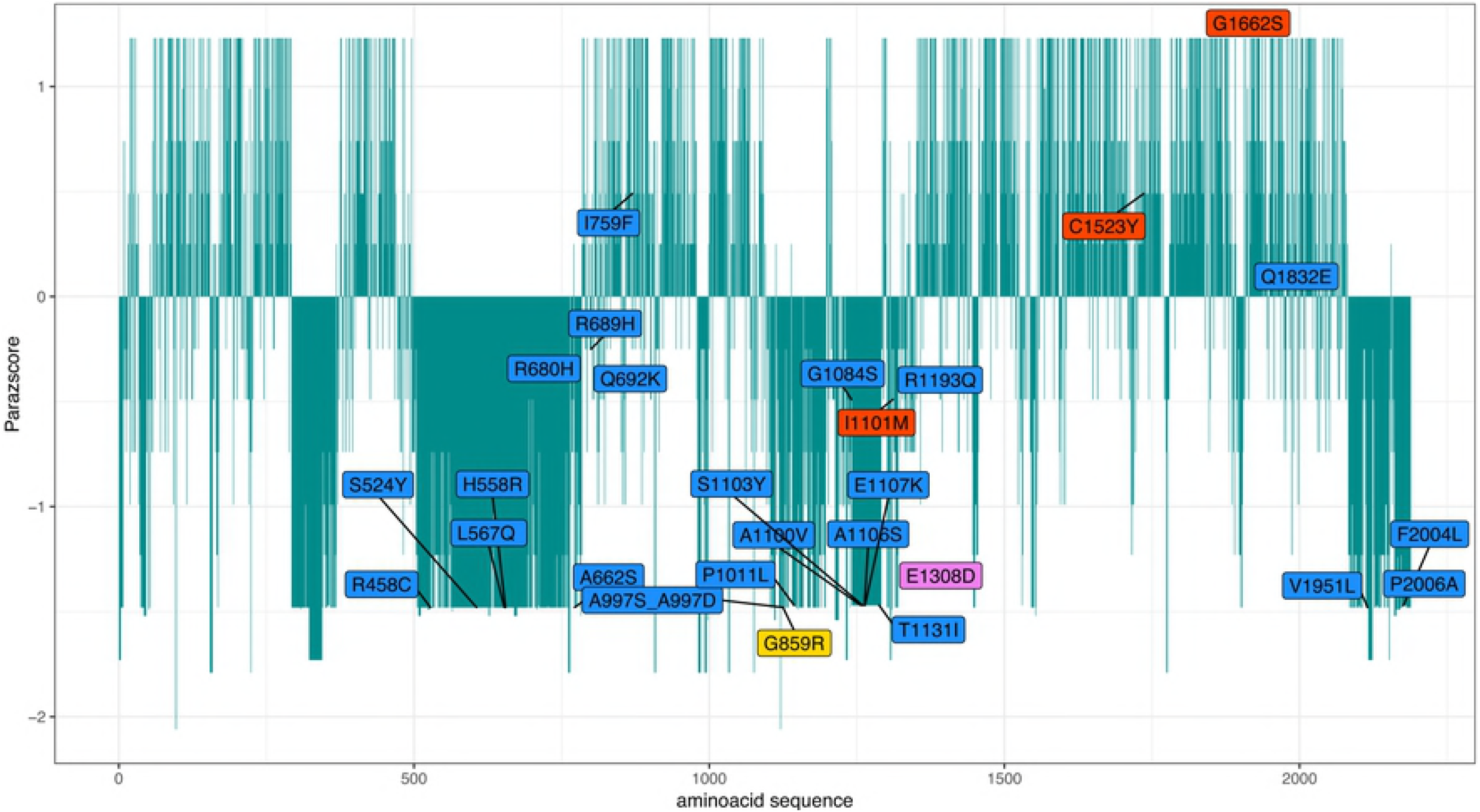
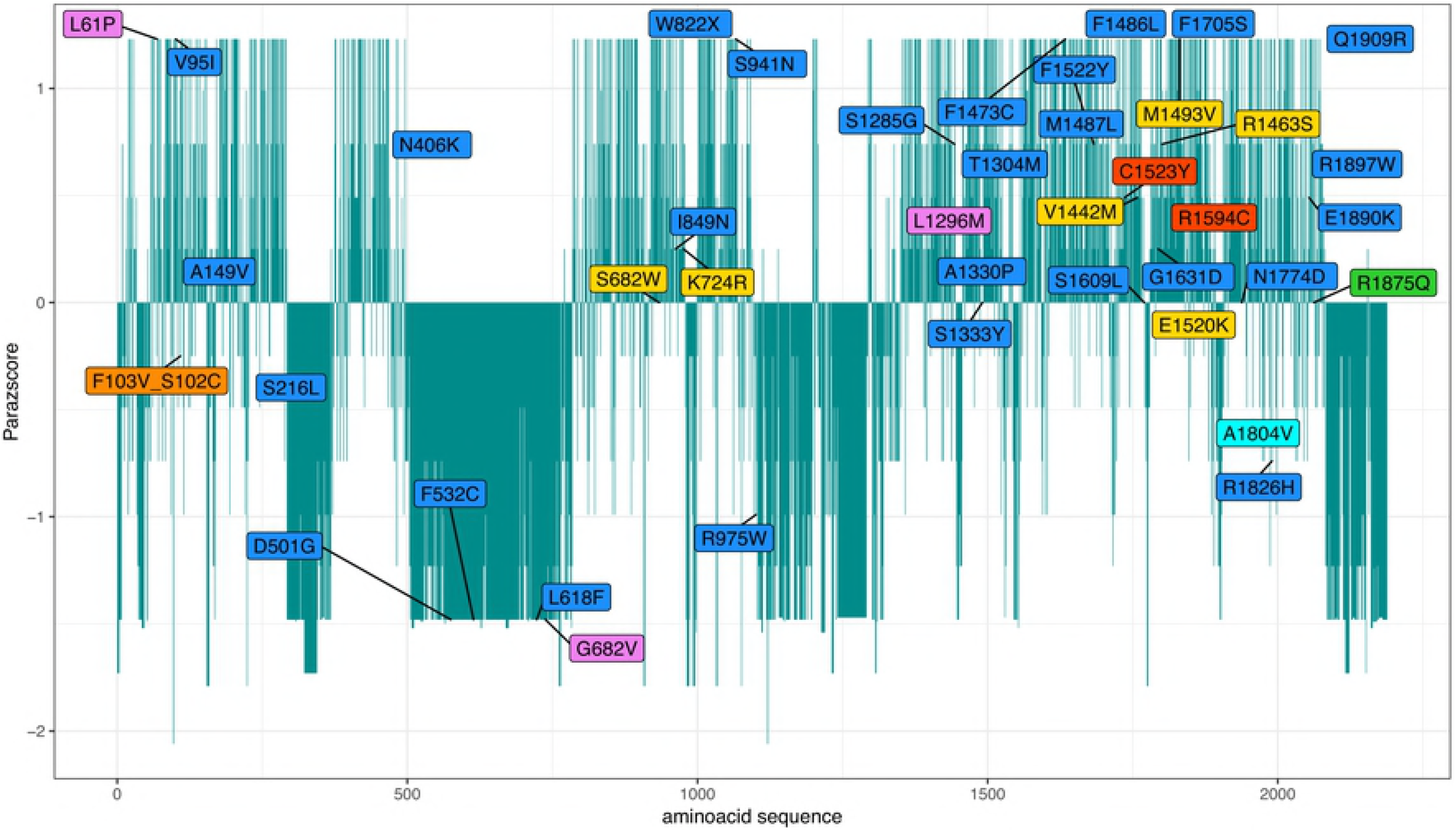
*SCN* patient variant evolutionary conservation and population constrained assessment. The *SCN* patient variant paralog conservation score (Parazscore) is shown across the linear protein sequence. Parazscore values range from negative values, representing less conservation at a given amino acid position, to positive values, representing high conservation, with the highest value depicting identical amino acids are present in all related proteins. A) The Parazscore is shown for *SCN* variants that are predicted to be pathogenic. B) The Parazscore is shown for *SCN* variants that are predicted to be conflicting. Color scale: purple = *SCN1A*, turquoise = *SCN3A*, yellow = *SCN4A*, orange = *SCN4A* and *SCN10A*, blue = *SCN5A*, green = *SCN9A*, red = *SCN10A*.

### Variant position and pathogenicity

Variants predicted to be pathogenic were more likely to be localized in the transmembrane regions of the protein (p-value = 0.03), which have been associated with severe channel dysfunction [45]. On the other hand, variants predicted to be conflicting or benign variants were more likely to be localized in the cytoplasm (p-value = 0.0063) (S1 Table, S1 Fig). Mapping of all Na_v_ channel variants on a 3-dimensional structural model was not informative (S2 Fig).

## Discussion

We report 11 patients with 13 pathogenic or likely pathogenic variants in genes encoding for sodium channel subunits expressed in the brain and/or in the heart. Among the cases with variants presented here, 8 cases had hippocampal abnormalities but no history of seizures. None had a history of cardiac arrhythmia or other cardiac presentation prior to death. While *SCN1A* [9,27], *SCN1B* [12,35-37], and *SCN5A* [12-26] have strong prior associations with epilepsy, arrhythmia, and sudden death, we additionally considered variants in the following *SCN* paralogs: *SCN3A, SCN4A, SCN9A, SCN10A*, Surprisingly, only 1/13 (8%) of our own cohort of patients carried a variant in *SCN5A*, compared to 76/99 (77%) of the SIDS patients reported in literature. On the other hand, 4/13 (31%) of our own cohort compared to 4/99 (4%) patients of the literature carried a variant in *SCN1A*, three of which were recently reported [9,27].

Our literature review found 74 variants in 99 patients with SIDS, SIDS-like presentations, or SUDC affecting 72 different amino acid positions in a VGSC-related gene. 55/75 variants were present in *SCN5A*, reflecting in some cases the fact that this gene was specifically targeted in some series [13-16,20,21,24,46]. Since many of the VGSC variants reported in SIDS or SUDC were described before the current era of abundant publicly available control data, we re-evaluated all the variants reported in literature with this in mind. In a surprisingly high number of reported variants (45%), conflicting evidence argued against pathogenicity using current ACMG criteria. Analysis of the relatively modest number of variants in our cohort, coupled with a larger number from the literature, suggested that those variants we determined to be pathogenic or likely pathogenic by other metrics were most likely to be located in critical domains of the sodium channel protein—e.g., transmembrane domains. Alternatively, variants in control populations were randomly distributed throughout the genes. Variants considered pathogenic are enriched in conserved regions across gene family members. We hypothesized that variants in VGSC genes at certain conserved positions might be associated with a risk for sudden death, independent of the tissue where the relevant gene is most highly expressed. This is illustrated by two patients in our cohort that carry a variant in respectively *SCN4A* and *SCN10A* at the same paralog position and 24 other variants from the literature that affect a paralog variant that is known to be disease-associated. When considering variant pathogenicity in a newly implicated VGSC-encoding gene, the position in the protein, with respect to paralogous proteins already implicated in sudden death, can provide additional evidence suggesting pathogenicity.

Ultimately, while all of these factors are taken into consideration when assessing pathogenicity, robustly conducted experimental evidence with adequate positive and negative control data should be sought when there is question regarding pathogenicity. Larger cohort studies will be required to more securely implicate a broader range of VGSC-related genes in SUDP. In addition, future studies that can incorporate trio sequencing, in which parental DNA can be made available to determine whether variants are *de novo* and thus more likely to be pathogenic with respect to a severe phenotype like sudden death, will contribute to our understanding of the role of this family of genes to sudden death. Initial studies in induced pluripotent stem cell (iPSC)-derived neurons and mouse models of *SCN1A*, traditionally associated with epilepsy, suggested cardiac and/or respiratory mechanisms of death [47,48]. Additional *in vitro* and *in vivo* studies of the sodium channel gene family will move us toward understanding the mechanisms through which variants in sodium channel-encoding genes contribute to sudden death.

## Conclusions

Our analysis of a SUDP cohort and the present literature on sodium channel variants in SUDP cases, using population- and protein structure-based predictive models, revealed 48 *SCN* variants. Importantly, in our cohort, the variants were found in children without prior histories of seizures yet with hippocampal abnormalities and without history or family history of cardiac arrhythmia. Overall, similar to those reported in prior studies, variants predicted to be pathogenic were more likely localized in the transmembrane regions of the protein. These findings provide evidence that sodium channel abnormalities contribute to the complex phenotype in SUDP involving central nervous system and/or cardiac rhythm dysfunction. Further, they suggest that future functional studies into the function of the sodium channels may elucidate the mechanisms through which variants in these genes underlie some cases of sudden death.

## Methods

### Ethics statement

The study has been approved by the Institutional Review Board of Boston Children’s Hospital (approval number P00011014). Informed consent has been obtained from all participants.

### Our cohort

DNA from 73 SUDP cases was obtained through the Massachusetts Office of the Chief Medical Examiner (OCME), Boston, MA and the Office of the Medical Examiner, San Diego, CA using consent procedures in accordance with Massachusetts and California Law. These cases included 42 singletons for whom parental samples were not available and for whom families could not be contacted, 28 trios consisting of probands and both parents, and 3 probands with one parent sample available. DNA extracted from whole blood or saliva underwent capture for exome sequencing using either the Agilent SureSelect XTHuman All Exon v4 or Illumina Rapid Capture Exome enrichment kit (Broad Institute, Cambridge, MA). Sequencing of 100bp paired end reads was obtained using Illumina HiSeq (Illumina, San Diego, CA). Coverage was >90% or >80% meeting 20x coverage with the two methods respectively. Our data analysis and variant calling methods have been described previously [49]. We utilized the BCH (Boston Children’s Hospital) Connect Genomics Gateway integrated with the *WuXi NextCODE* analysis platform [50] for variant interrogation and analysis.

For each case, we performed a targeted initial analysis to identify variants in genes encoding for the human VGSC subunits. Candidate pathogenic variants were evaluated according to American College of Medical Genetics and Genomics (ACMG) criteria [44], including pathogenicity predictions from both Polyphen2 and SIFT and low population allele frequency (<0.001) according to the Genome Aggregation Database (gnomAD, http://gnomad.broadinstitute.org). For cases with data from parental samples, we evaluated *de novo* vs. inherited status of candidate variants of interest. For splicing variants, we used the splicing prediction score from *Alamut Visual-2.10*, which incorporates the splicing tools MaxEnt, NNSPLICE, and HSF.

### Literature cohort

In order to identify additional cases for phenotypic comparison and to evaluate whether a given variant was novel or previously reported and whether there might be data supporting pathogenicity, we performed a literature search (PubMed, accessed June 2018, with search parameters “Sudden Infant Death” [Mesh] AND “Sodium Channels” [Mesh] resulting in the identification of 43 studies. In addition, we searched the Human Gene Mutation Database (http://www.hgmd.cf.ac.uk/, accessed June 2018) for each of the VGSCs genes to identify any possible variants not found in the literature search, identifying 13 additional studies with cases of SIDS or SUDC and reported variants in sodium channel-related genes.

### Structural protein modeling

We assessed genotype-phenotype correlation by comparing the locations of variants and the phenotypic features associated with each variant using the human Na_v_1.7 (*SCN9A*) protein model described by Huang et al [51]. We analyzed the position of the variants of our cohort along with additional variants identified through our literature search (S1 Table). Three of our cases’ variants have been previously reported in literature [9,27]. We focused on exonic variants since intronic, splicing, and truncating variants cannot be annotated onto the three-dimensional protein sequence. Illustrations were generated using PyMol.

### In silico predictions

Functional prediction scores were obtained from the dbNSFP database version 3.5 (August 2017, http://varianttools.sourceforge.net/). In total, we used six pathogenicity prediction scores (SIFT, Polyphen-2-HVAR, Polyphen-2-HDIV, Mutation Assessor, FATHMM, and LRT). We classified a variant as “damaging” when the majority of the tools predicted a functional effect for the variant (i.e., a minimum of 4 out of 6 tools). Splicing variants are considered “possibly damaging” or “damaging” when they have a likelihood of 50% or more to affect splicing. For splicing variants, we used the splicing prediction score from Alamut Visual-2.10, which is calculated from the splicing tools MaxEnt, NNSPLICE, and HSF.

### Parazscore

Based on the linear amino acid sequence of *SCN9A* (canonical transcript ENST00000409672, CCDS46441), we compared the position of 74 missense variants in all sodium channel gene paralogs against variants found in the general population using gnomAD. We evaluated the amino acid gene-family paralog conservation score using the Parazscore [52] (http://mbv.broadinstitute.org), which leverages amino acid conservation across gene-family members, assuming that conserved sites are more likely to be important for protein function and thus more likely to be present in patients than in controls. Statistical comparison between the variant counts of patients vs. gnomAD was conducted using a two-tailed t-test with nominal two-sided p-values <0.05 considered significant.

### Three-dimensional mapping of amino acid substitutions in VGSC-related genes

In order to assess for a genotype-phenotype correlation across gene-family paralogs, we compared the position of all missense variants from our own cohort and from literature onto a 3-dimensional Na_v_1.7 structure model. Since no atomic structure of any mammalian Na_v_ channel is available, we used the recently published Na_v_1.7 structure model [51] that has been established on the cryo-EM structure of a rabbit Cav channel Cav1.1.

## Supporting information

**S1 Fig. Pathogenic *SCN* patient variants are more likely to be localized in the transmembrane regions.** The distribution of the paralogues of the *SCN* patient variants is shown on a two-dimensional protein model of the *SCN9A* protein. Variants predicted to be pathogenic are shown in red. Variants predicted to be benign or conflicting are shown in green.

**S2 Fig. *SCN* patient variant distribution on a three-dimensional protein model.**

The distribution of the paralogues of the *SCN* patient variants is shown on a threedimensional protein model of the *SCN9A* protein, described by Huang et al[51]. The backbone of the protein is shown in white, and disease-associated variants are shown as red spheres.

**S1 Table. All variants in VGSC genes in patients with SIDS or SUDC**

Legend: ‘patient has two variants in *SCN5A;* *patient has two variants in *SCN5A*, ◆ carries a variant *in SCN5A* and *SCN1B* (both inherited from father), ○ compound heterozygous for *SCN5A*, □ compound heterozygous for *SCN1A*. AF = allele frequency; D = deleterious; gnomAD = Genome Aggregation Database; GOF = gain of function; H = high; n patients = number of patients; L = low; LOF = loss of function; LRT = likelihood ratio test; M = medium; MA = Mutation Assessor; N = neutral; P = pathogenic; PP-2 = PolyPhen2; SIFT = Sorting Intolerant from Tolerant; T = tolerant. Transcripts used: *SCN1A* (NM_001165963.1), *SCN3A* (NM_0006922.3), *SCN4A* (NM_000334.3), *SCN5A* (NM_198056.2), *SCN9A* (NM_002977.3), *SCN10A* (NM_006514.2), *SCN1B* (NM_199037.4), *SCN3B* (NM_018400.3), *SCN4B* (NM_174934.3).

**S2 Table. Patients with variants at a paralog position**

Legend: AF = allele frequency; GOF = gain of function; LOF = loss of function; N/A = not available. Variants in bold were identified in our cohort. Transcripts used: *SCN1A* (NM_001165963.1), *SCN3A* (NM_0006922.3), *SCN4A* (NM_000334.3), *SCN5A* (NM_198056.2), *SCN9A* (NM_002977.3), *SCN10A* (NM_006514.2), *SCN1B* (NM_199037.4), *SCN3B* (NM_018400.3), *SCN4B* (NM_174934.3).

## Acknowledgments

The authors are grateful to all of the families who participated and continue to participate in research on SIDS, SUDC, and SUDEP. We are deeply indebted to Dr. Hannah C. Kinney and her dedication to Robert’s Program; her neuropathological analysis of SIDS and SUDC patient samples provided a major contribution to the present study. We thank Kalen Fletcher for facilitating patient enrollment at Robert’s Program at Boston Children’s Hospital.

